# CD8 T cell hyperfunction and reduced tumour control in models of advanced liver fibrosis

**DOI:** 10.1101/2023.09.20.557752

**Authors:** Jood Madani, Jiafeng Li, Agatha Vranjkovic, Katrina Jorritsma, Mohamed S Hasim, Manijeh Daneshmand, Angela C Cheung, Angela M E Ching, Jennifer E Bruin, Michele Ardolino, Angela M Crawley

## Abstract

Immune dysfunction, both depression and hyperactivation, in liver disease contributes to significant morbidities and mortalities, depending on liver damage severity and etiology. The underlying causes of immune dysfunction in advanced liver disease, whether pathogen or host-mediated, remain unclear. We reported lasting generalized CD8^+^ T cell hyperfunction in individuals with advanced liver fibrosis in chronic HCV infection. The separation of viral and fibrosis-driven effects or the association of this phenomenon with clinical outcomes of advanced liver fibrosis remains to be determined. Here, a hepatotoxic murine model of liver fibrosis was used to decouple liver fibrosis from viral infection. Carbon tetrachloride (CCl_4_)-treated mice presented progressive liver fibrosis within ≈12 weeks, resulting in severe diffuse fibrosis, focal necrosis and surrounding mixed inflammation; pathology similar to that of chronic HCV infection. Taking advantage of this model, we investigated if liver fibrosis caused systemic CD8^+^ T cell hyperfunction and evaluated its impact on host immune response. At peak liver fibrosis, circulating CD8^+^ T cells presented increased expression of IFN-γ and granzyme B (GrzB) in comparison to control animals. CD8^+^ T cell hyperfunction arose by 8 weeks of CCl_4_ treatment and was sustained with continued liver insult. As a result, fibrotic mice were unable to resist an ectopic tumour challenge and were less responsive to immunotherapy. Furthermore, CD8^+^ T cell dysfunction was observed in other contexts of chronic liver insult such as high fat diet-induced liver steatosis, even in the absence of significant fibrosis. Collectively, this study shows the impact of chronic liver insult on systemic CD8^+^ T cell function and its association with impaired immune response, such as tumour surveillance.

## Introduction

Increasing evidence shows that the liver is an important immunological organ, wherein immune cells reside and transition to be educated as well as routinely traffic with blood flow. Therefore, unsurprisingly, liver disease is also associated with major acquired defects in liver and systemic immune function^1,2^. An estimated 10-25% of the global population, including 1 in 4 Canadians, are affected by recent surges in the incidence of chronic liver disease, caused predominantly by non-alcoholic fatty liver disease (NAFLD), alcohol use disorder (AUD), the hepatitis C virus (HCV) epidemic largely attributable to intravenous drug use, as well as viral hepatitis in endemic immigrant communities^3^. Collectively, these diseases contribute to rising deaths from cirrhosis and the doubling of hepatocellular carcinoma (HCC) deaths.

While the underlying cause of fibrosis can be eliminated, allowing hepatic tissue remodelling (e.g. direct-acting antiviral (DAA)-mediated HCV clearance, dietary intervention in NAFLD and alcohol abstinence in AUD), whether immune dysfunctions are reversible is unknown. Chronic HCV infection disrupts immune function, affecting many innate and adaptive immune cells including cytotoxic CD8^+^ T cells^4, 5, 6, 7, 8, 9^. Despite the spectacular results of DAAs, it is not clear if sustained virologic response (SVR) parallels restored immune function. There are contrasting reports in the literature, many of which demonstrate some recovery of previously exhausted HCV-specific T cells^10, 11, 12, 13, 14, 15^, although this is not always the case in those with advanced liver fibrosis^16, 17, 18^.

We, and others, have observed significant impairment of the entire CD8^+^ T cell compartment in the blood and liver of HCV patients^12, 13, 14, 15^ and decreased expression of pro-survival Bcl-2 in CD8^+^ T cells from patients with advanced liver fibrosis^19^. We subsequently detected an increased function profile in blood-circulating CD8^+^ T cells of HCV-infected individuals with cirrhosis compared to those with minimal fibrosis, which persisted long after achieving SVR with DAA therapy^20^. The molecular and cellular mechanisms linking liver fibrosis severity in chronic liver disease to such immune dysfunction, and the consequences thereof, are not clear. It remains important to better understand this, as restoring normal function *in vitro* in HCV/CMV/EBV-specific CD8^+^ T cells from HCV-infected individuals with advanced liver fibrosis remains a challenge^21^.

Much has been learned about liver fibrosis progression in mouse models^22^, which are valuable as complementary models of chronic HCV infection with significant liver damage. While there is no single animal model that recapitulates all the features of liver disease, the carbon tetrachloride (CCl_4_) model is characterized by a linear development of liver fibrosis followed by periportal and portal-to-central fibrosis, both hallmark features of HCV infection pathology^22^. This hepatotoxin model is a widely used and reliable animal model of progressive hepatic fibrosis as well as fibrosis regression^22, 23, 24^. Furthermore, the CCl_4_ model uncouples liver fibrosis from the confounding factors derived from a hepatic virus infection, such that any effect observed on the immune system can be attributed to the former and not to the latter. We hypothesize that associating liver fibrosis severity to immune dysfunction is of broader relevance to liver disease regardless of etiology.

In this study, we characterized and examined the functional disruptions of peripheral CD8^+^ T cells in the hepatotoxin model. Animals administered CCl_4_ intraperitoneally (i.p.) for approximately 12 weeks consistently developed significant chronic liver fibrosis^23^. The responses of circulating CD8^+^ T cells were overactive in mice with advanced liver fibrosis, corroborating our hypothesis that liver fibrosis is sufficient to trigger immune dysfunction. Furthermore, we complement these findings with data showing CD8^+^ T cell hyperfunction in a model of high fat diet-induced steatosis with minimal fibrosis, broadening this immune dysfunction to different modalities of chronic liver disease.

## Methods

### Study approval and animals

Mouse studies were performed in accordance with the guidelines of the Canadian Institutes of Health Research. For the CCl_4_ experiments, C57BL/6 mice were purchased from the Jackson Laboratory and housed at the University of Ottawa Animal Care Veterinary Services (ACVS) facilities, certified by Canadian Council on Animal Care. This experimental protocol was reviewed and approved by ACVS at the University of Ottawa. Across 6 independent studies, we used a total of 87 females and 51 males, ranging between 7-12 weeks old at experimental onset. The high fat diet experiments included male and female C57BL/6 mice, which were bred and housed in the Carleton University vivarium. The latter study was approved by the Carleton University Animal Care Committee and carried out in accordance with the Canadian Council on Animal Care guidelines.

### CCl_4_ induction of advanced liver fibrosis

To achieve a state of advanced liver fibrosis, C57BL/6 mice were injected twice weekly with CCl_4_ (1.0 ml/kg, Sigma-Aldrich, USA) in olive oil (Bertolli), as an adaptation of established protocols^25,26^. At predetermined time points, animals were euthanized, and livers were immediately harvested, formalin-fixed and subsequently embedded in paraffin. To evaluate fibrosis severity, liver tissue sections were stained with Masson’s trichrome and scored by a skilled pathologist (Nour Histopathology Consultation Services, Ottawa, ON, Canada) following the METAVIR system (F0-1 minimal fibrosis to F4 advanced fibrosis/cirrhosis), which is used both clinically and in rodents^27^. Liver fibrosis regression was evaluated following the cessation of CCl_4_ injections after ≈ 12 weeks, alongside continued control/CCl_4_ treatments for a minimum of 4 weeks.

### Cell isolation and culture

Blood samples were collected by saphenous vein draw and lysed with Hybri-Max™ red blood cell lysing buffer (Milipore-Sigma). Resulting PBMCs were cultured at 1×10^6^ cells/mL in 96-well high binding plates (Sarstedt, USA) pre-coated for 1 hour at 37°C with anti-CD3 antibodies (5ug/mL, clone: 145-2C11, BD Biosciences, San Diego, CA, USA) and supplemented with soluble anti-CD28 antibodies (2ug/mL clone: 37.51, BD Biosciences). Cells were cultured for 48 hrs before analysis.

### Flow cytometry analysis of CD8^+^ T cell phenotype and function

Six hours prior to the end of the 48-hour anti-CD3/-CD28 cell stimulation, cells were treated with Golgi Plug (BD Biosciences) and anti-mouse CD107a antibody was added. Next, live/dead cell staining was added (Live-Dead Fixable Stain Kit, Molecular Probes, Eugene, Oregon, USA). Cells were then labelled with cell surface receptor antibodies using the following antibodies: anti-CD3 (clone: UCHT1, BV-510), CD4 (clone: GK1-5, AlexaFluor 700), CD8α (clone: 53-6.7, BV-785), CD19 (clone: 6D5, Pe-Cy5), CD44 (clone: IM7, BV-421), CD62L (clone: MEL-14, PE), PD-1 (clone: 29F.1A12, PE-Cy7), CD107a (clone: 1D4B, APC) from BioLegend (San Diego, CA, USA), and NKG2D (clone: CX5, BV-711) BD Biosciences (San Diego, CA, USA). Cells were then fixed and permeabilized with CytoFix/CytoPerm intracellular staining kit (BD Biosciences) and labelled with anti-mouse TNF-ɑ (clone: Mab11, FITC), IFN-γ (clone: 4S.B3, BV-650) (BioLegend, San Diego, CA, USA), and GrzB (clone: GB11, Texas Red) (Horizon, BD Biosciences, San Diego, CA, USA). The following subsets were distinguished in this study: naïve (T_N_, CD45RA^+^CD62L^+^), effector (T_E_, CD45RA^-^CD62L^-^) and central memory (T_CM_, CD45RA^-^CD62L^+^). Cells were then evaluated by flow cytometry using an LSR-Fortessa (BD, San Diego, CA). Acquired flow cytometry data was analyzed using FlowJo software (FLOWJO, LLC, Ashland, Oregon).

### Cancer challenge in chronic liver fibrosis

To investigate the effects of liver fibrosis on responses to tumour challenge, in some experiments, mice were injected subcutaneously (S.C.) with MC38 tumour cells (1×10^5^ – 1×10^6^ cells in 100 µL PBS). Once palpable, ectopic tumour growth was measured every two days using calipers. For immunotherapy experiments, mice were challenged with MC38 tumour cells and once tumours reached ~50 mm^3^, they were treated with anti-PD-1 anti-CTLA-4 antibodies (clones RMP1-14 and 9H10, respectively, from Leinco Technologies Inc., Fenton, MO, USA): 200 µg ab/injection, i.p. in 100 µL PBS, 3 injections, 2 days apart.

### High-fat diet induced liver disease

To induce liver disease in a high-fat diet (HFD) model, a 14-week study was conducted. Mice were placed on a regular chow diet (Teklad Diet #2014 –13% fat, 20% protein, 67% carbohydrates) after weaning. At 28 to 30 weeks of age, a subset of mice (6 males, 8 females) was transferred to a 45% high-fat diet (Research Diets, D12451) for a duration of 14 weeks, while the control group (8 males, 8 females) remained on the regular chow diet. Body weight was measured weekly, and fat and lean mass were subsequently measured relative to total body weight at week 11 using an EchoMRI-700 (EchoMRI LLC, Houston, TX, USA), A glucose tolerance test was performed after 13 weeks to assess systemic glucose homeostasis. At endpoint, liver tissue was harvested, stored in 4% paraformaldehyde, transferred to 70% ethanol until paraffin embedding and sectioned to evaluate liver fibrosis, steatosis, ballooning, and inflammation by a pathologist (Nour Histopathology Consultation Services, Ottawa, ON, Canada).

### Data analysis

Data were analyzed by either two-tailed one-way Students’ *t*-test, one-way ANOVA with Dunnett’s post-test or correlation analyses (p≤0.05), as appropriate. GraphPad Prism 5.0-10.0.1 software was used for statistical analyses and to plot data.

## Results

### CCl_4_-treated mice develop advanced liver fibrosis mirroring pathologies observed in human HCV^+^ patients

To examine the effect of developing advanced liver fibrosis on immune function, in the absence of viral infection or other confounders, a mouse model of chronic liver fibrosis induced by CCl_4_ was adapted from established protocols^23, 25, 26^. Mice were injected twice weekly with CCl_4_ (1 ml/kg, i.p.) for 10-14 weeks, as described in Figure 1A. We determined that this dose was well tolerated by carefully monitoring animal vital signs and weight gain, which remained relatively similar between groups (Supplemental Figure 1). For histological analyses, livers were harvested from randomly selected mice at varying treatment timepoints.

**Figure 1:**
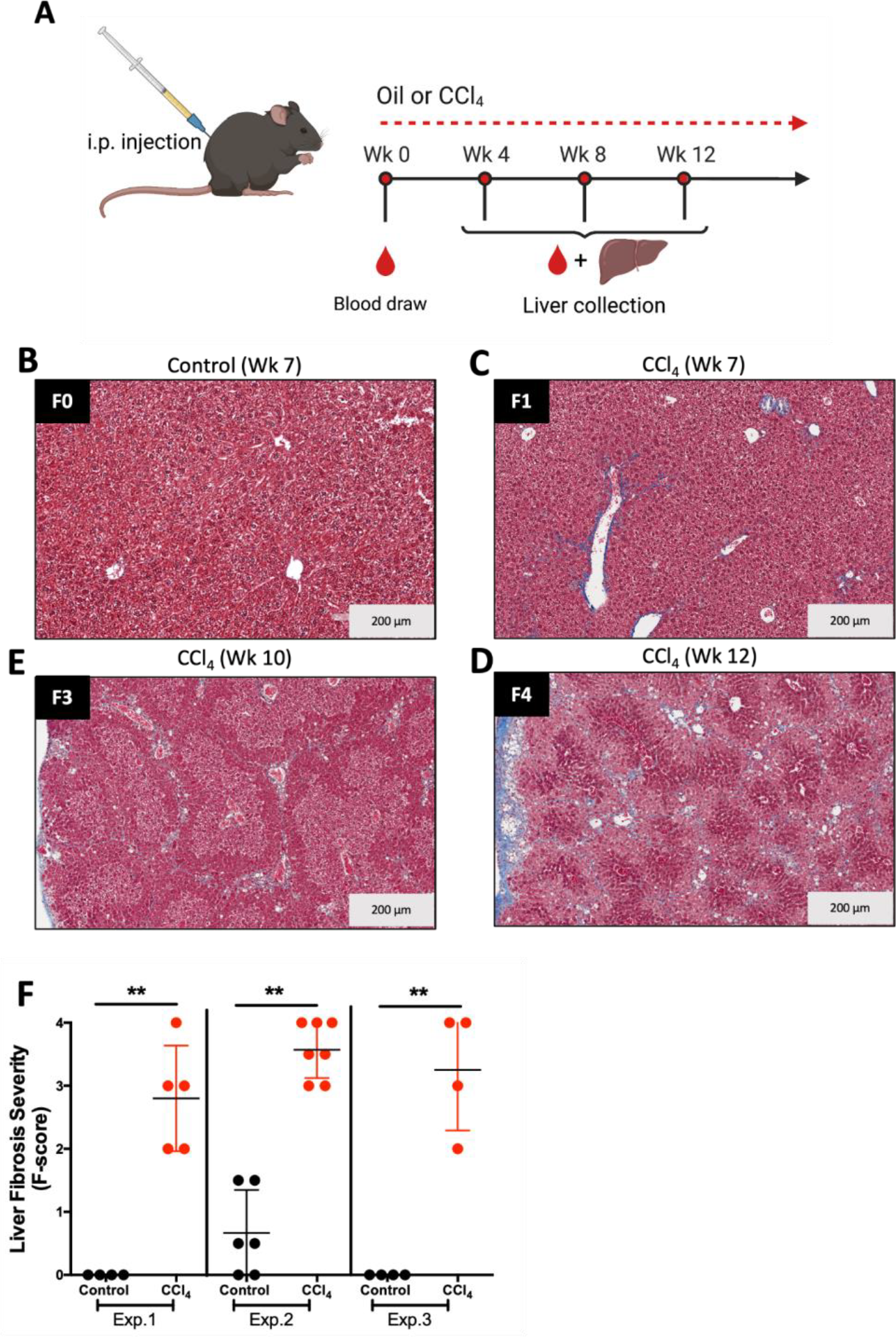
CCl_4_-induced liver fibrosis. **A)** A schematic of the experimental design of CCl_4_ administration in C57BL/6 mice. Mice were injected i.p. with oil (control) or CCl_4_ in oil (1ml/kg body weight) twice/week for 12-16 weeks. Blood was collected every 4 weeks to assess circulating CD8^+^ T cell phenotype and function. At various time points, livers were harvested from euthanized mice. **B-D)** Representative tissue images demonstrate liver fibrosis severity. Livers were formalin-fixed, paraffin embedded, sectioned and stained with Masson’s Trichrome three-colour stain. Fibrosis staging was determined by a pathologist using the METAVIR (F-score) scoring system: F0 = no fibrosis, F1/2 = minimal liver fibrosis, F3/4 = advanced fibrosis. Images represent scanning microscopy at 10x magnification, for: **B)** Control after 7 weeks of oil i.p., and CCl_4_-treated mice after **C)** 7 weeks; **D)** 10 weeks; and **E)** 12 weeks. **F)** A summary of liver fibrosis scores from 3 independent experiments. Data are presented as means ±SD, and statistically significant differences were determined using a two-tailed unpaired Students’ *t*-test (**p<0.001).

After 7 weeks, liver tissue in control mice remained healthy (Figure 1B), whereas those injected with CCl_4_ began to develop moderate peripheral and bridging fibrosis (F1, Figure 1C). By week 10, liver tissue in CCl_4_-treated mice progressed to severe diffuse fibrosis, focal necrosis and surrounding mixed inflammation (F3-F4, Figures 1D and E). Across several independent experimental groups, this 10–14-week CCl_4_ administration protocol resulted in the consistent induction of advanced liver fibrosis (Figure 1F). The fibrosis patterning resembles that of chronic HCV infection, with characteristic bridging fibrosis pathology leading to cirrhosis^22^. With this model, we determined there to be no significant sex differences in CCl_4_-induced liver fibrosis. These findings highlight the validity of the hepatotoxin model to study the effect of progressive liver fibrosis on the immune system decoupled from viral infection.

### Mice with advanced liver fibrosis exhibit generalized CD8^+^ T cell hyperfunction

To determine if CD8^+^ T cell hyperfunction is associated with liver fibrosis severity in this animal model, as previously observed in cirrhotic HCV-infected patients^28^, the functionality of peripheral blood CD8^+^ T cells was assessed at peak of liver fibrosis (i.e. ≈12 weeks CCl_4_ treatment). The proportion of GrzB^+^CD8^+^ T cells from CCl_4_-treated mice was significantly higher than in control mice (Figure 2A). This result was replicated in several independent experiments (data not shown). No significant differences were detected in the CD107a degranulation marker between groups (Supplemental Figure 2A).

**Figure 2:**
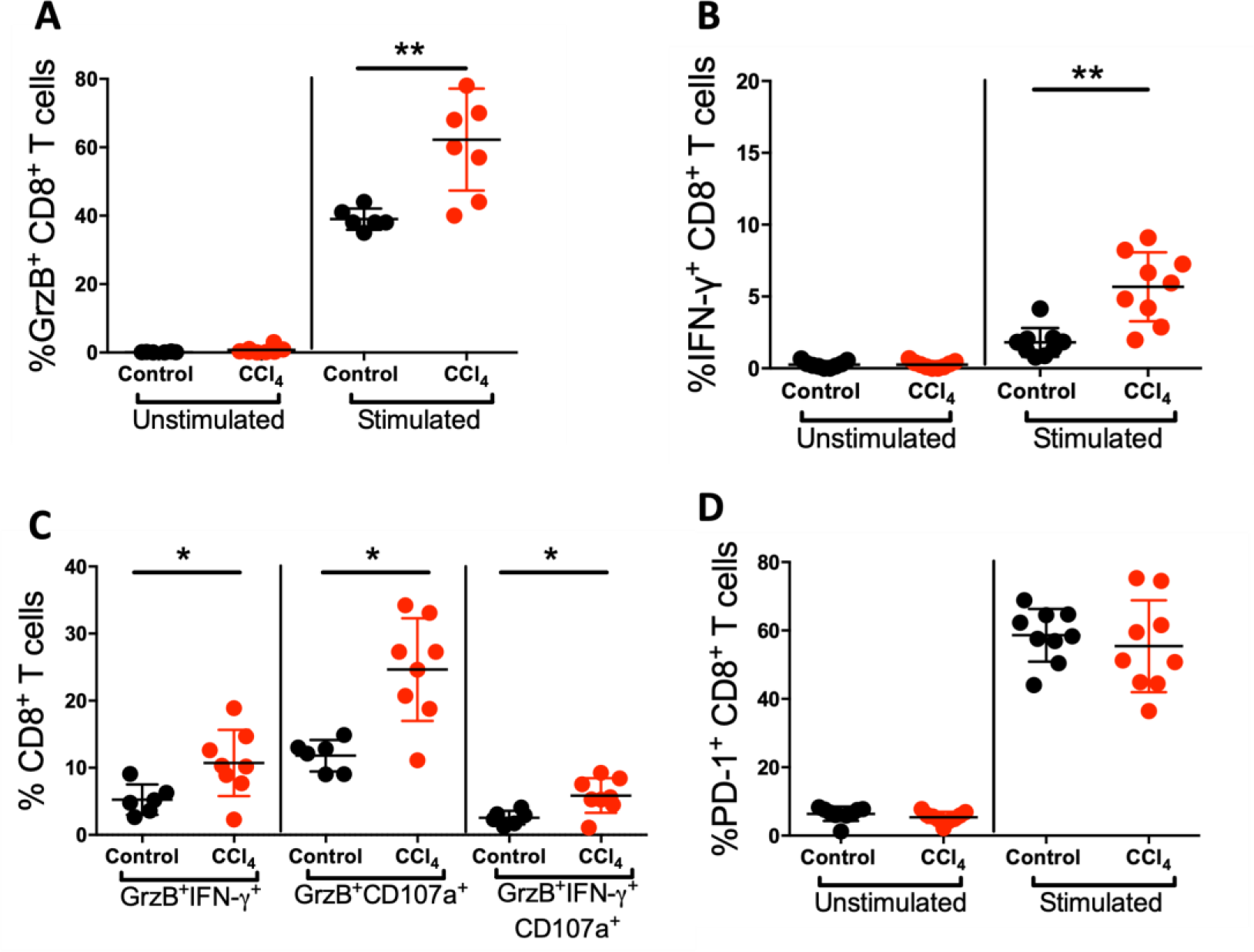
Systemic CD8^+^ T cell hyperfunction is associated with advanced liver fibrosis. Isolated PBMCs were stimulated *in vitro* with anti-CD3/28 antibodies for 48 hours, and function was assessed by flow cytometry. Summary data of **A)** GrzB and **B)** IFN-γ expression in control and CCl_4_-treated mice after 12 weeks of the treatment (latter mice were confirmed histologically to have developed advanced liver fibrosis). **C)** Increased proportions of polyfunctional (i.e., IFN-γ^+^GrzB^+^, CD107a^+^GrzB^+^) and oligo-functional (CD107a^+^ GrzB^+^ IFN-γ^+^) CD8^+^ T cells in CCl_4_-treated mice compared to controls. **D)** The % expression of activation marker PD-1 on CD8^+^ T cells in both study groups. Data are presented as means ±SD and statistical significance was determined using a two-tailed unpaired Students’ *t*-test (*p<0.05, or **p<0.001).

In CCl_4_-treated animals, the expression of immune-modulating IFN-γ was assessed and found to be significantly elevated, by two-fold, compared to control animals (Figure 2B). There was also a significant elevation in the proportion of bi-functional IFN-γ ^+^GrzB^+^ CD8^+^ T cells in treated animals (Figure 2C,). Similarly, there were two-fold more CD107a^+^GrzB^+^ cells in the treatment group compared to controls (Figure 2C). Furthermore, co-expression of all three analytes was also elevated in CCl_4_-treated mice. Taken together, the data suggests that both cytokine production and the cytotoxic potential of bulk circulating CD8^+^ T cells are altered in mice with advanced liver fibrosis, individually and concurrently. It should be noted that CD8^+^ T cells from both study groups appeared to be similarly activated in response to stimulation, as measured by an equivalent upregulation of PD-1 expression (Figure 2D).

In our previous study, the observed generalized CD8^+^ T cell hyperfunction in HCV-infected individuals with advanced fibrosis was pronounced in naïve (T_N_), central (T_CM_) and effector memory (T_E_) cell subsets, particularly for IFN-γ or perforin expression^28^. Therefore, a subset-specific analysis was conducted in the CCl_4_ mouse model and dynamic expression patterns were observed in a multi-parameter flow cytometry analysis. In the assessment of cytolytic potential, we found elevated proportions of GrzB^+^ cells in all CD8^+^ T cell subsets from CCl_4_-treated mice compared to controls (Figure 3A). Furthermore, we observed that the T_E_ cell subset exhibited the highest proportion of CD107a^+^ cells after stimulation, followed by T_CM_ and T_N_ cells. However, in bulk CD8^+^ T cells, there were no significant differences in the proportion of CD107a^+^ T cell subsets, except for an increase in the T_CM_ subset, following stimulation between the treatment group and controls (Supplemental Figure 2B).

**Figure 3:**
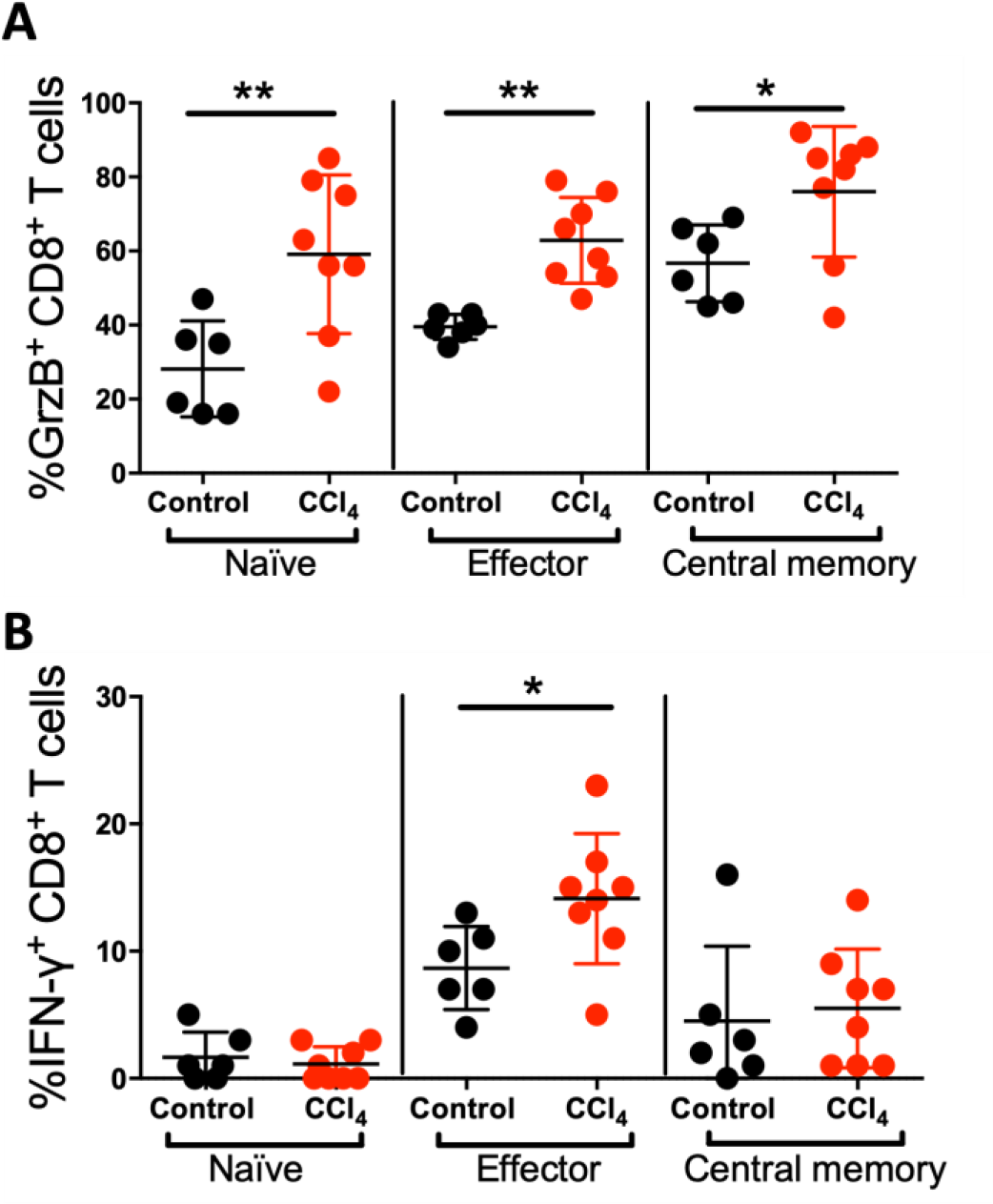
CD8^+^ T cell hyperfunction across T cell subsets. Isolated PBMCs were stimulated *in vitro* with anti-CD3/28 for 48 hours. T cell subset function was assessed by flow cytometry using subset distinguishing phenotypes: naïve (CD44^low^CD62L^+^), effector (CD44^+^CD62L^-^), and memory (CD44^+^CD62L^+^). Significant increase in the proportion of **A)** GrzB^+^ and **B)** IFN-*y* ^+^ cells in all CD8^+^ T cell subsets in mice treated with CCl_4_ for 12 weeks with advanced liver fibrosis compared to controls. Data are presented as means ±SD and statistical significance was determined using a two-tailed unpaired Students’ *t*-test (*p<0.05, or **p<0.001).

As with GrzB, the highest proportion of IFN-γ^+^ cells after stimulation were found in the T_E_ cell subset, followed by T_CM_ and T_N_ cells in both study groups. Specifically, an increased proportion of IFN-γ ^+^ T_E_ CD8^+^ T cells was observed in CCl_4_-treated mice in comparison to controls (Figure 3B). These findings were confirmed in three subsequent experiments. In summary, CD8^+^ T cell hyperfunction in this animal model of liver fibrosis is marked by elevated IFN-γ expression, largely driven by the T_E_ cell subset, and bears remarkable similarity to observations in humans with cirrhosis in HCV infection^28^.

### Mice with cirrhosis exhibit sustained CD8^+^ T cell hyperfunction despite liver fibrosis regression

In humans, the removal of chronic liver insult, such as hepatic viral infection, can result in the regression of liver fibrosis, albeit dependent on the initial severity of liver damage^29, 30^. It has been previously reported that CCl_4_-induced liver fibrosis in rats is reversed 28 days (i.e., 4 weeks) post-treatment cessation, characterized by a progressive loss of fibrotic matrix^31^. To determine if liver fibrosis regression will occur in a mouse model, treatment was stopped in a subgroup of mice after 12 weeks of hepatotoxin administration, followed by liver histology 4 weeks later (Figure 4A). By week 16, liver fibrosis severity reversed significantly from F3-4 to F1 in most CCl_4_-stopped mice, while fibrosis persisted in treatment-continued mice, as expected. (Figure 4B-E).

**Figure 4:**
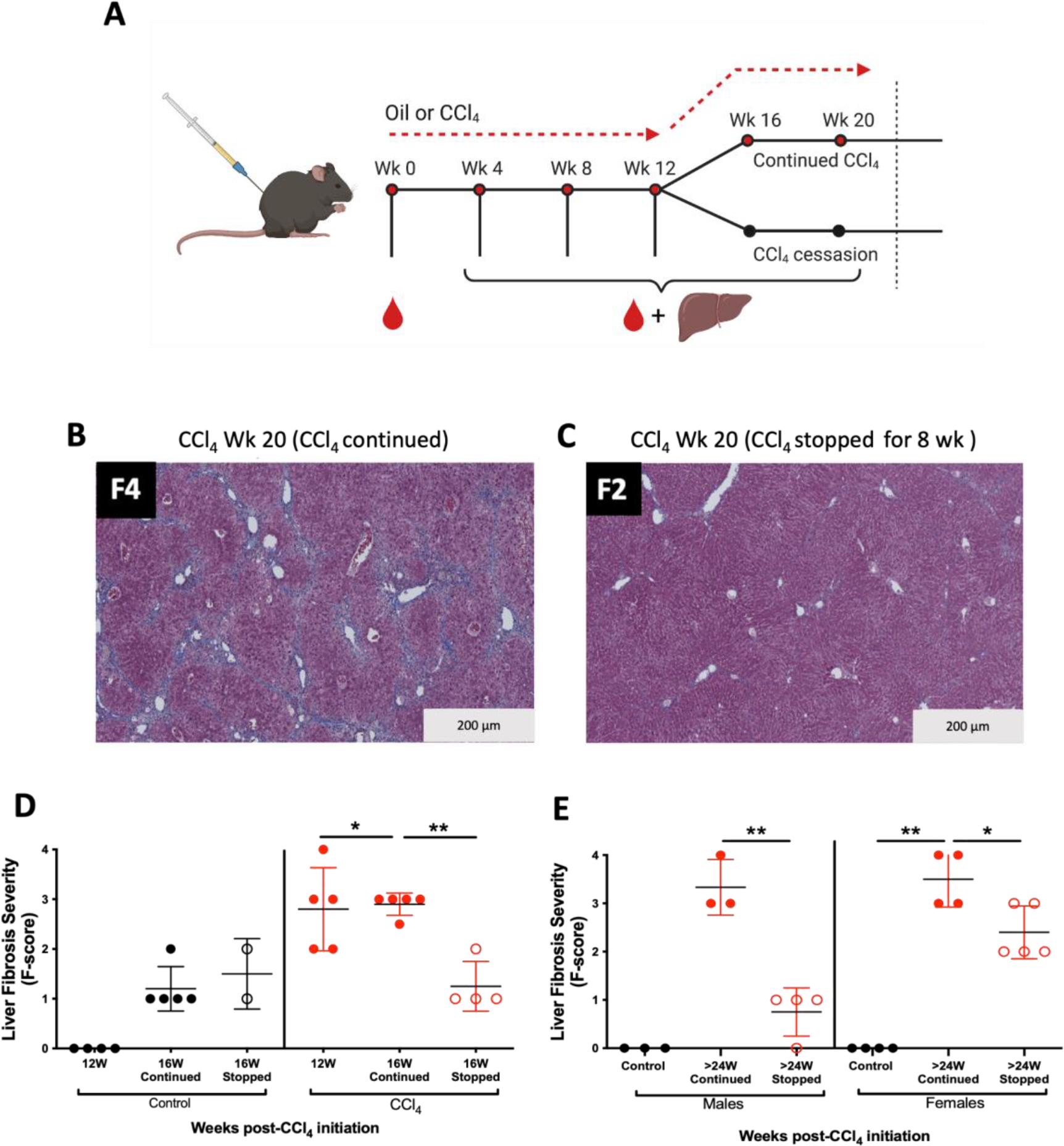
Liver fibrosis regression post-CCl_4_ cessation. **A)** A schematic representing the experimental design for the continuation or cessation of CCl_4_ treatment after 12 weeks (i.e., after peak fibrosis is achieved). Mice were injected i.p. twice weekly with oil (control) or CCl_4_ in oil for up to 12 weeks. At this time, CCl_4_-treated mice were randomly divided into two groups: 1) continued CCl_4_ administration for up to 20 weeks (CCl_4_ continued), and 2) CCl_4_ stopped, to allow for liver fibrosis regression. Representative liver histology images of Masson’s Trichrome three-colour staining show liver fibrosis severity with **B)** continued CCl_4_ for 20 weeks and **C)** 8 weeks after cessation of CCl_4_ treatment; fibrosis scores are indicated (10x magnification). **D)** A summary of liver fibrosis severity for mice with 20 weeks of continued CCl_4_ injections or cessation for 4 weeks after initial 12-week regimen of injections. **E)** Liver fibrosis progression and regression in all treatment groups with continued CCl_4_ treatment, or stopped oil/CCl_4_ treatment, between males and females. Data are presented as mean METAVIR score ±SD and statistical significance was determined using a two-tailed unpaired Students’ *t*-test (*p<0.05 or **p<0.01).

We previously reported that CD8^+^ T cell hyperfunction in HCV^+^ individuals persisted 12 months after the initiation of a successful course of DAA therapy, regardless of the degree of liver fibrosis regression. In those with advanced liver fibrosis, regression was minimal^28^. As shown, mice with hepatotoxin-induced liver fibrosis can undergo partial or full regression 4 weeks after the cessation of CCl_4_ administration. Following cessation, we found sustained detection of elevated CD107a and IFN-γ expression, similar to those continuously administered CCl_4_ (Figure 5A, B). In contrast to these findings, there was no difference in the proportion of GrzB^+^CD8^+^ T cells in control and CCl_4_-experienced groups after treatment cessation. This may have been driven largely by the noted significant decrease in GrzB^+^CD8^+^ T cells in female mice with CCl_4_ cessation whereas proportions of GrzB^+^ cells in males were equivalent to controls (Figure 5C). Therefore, after the removal of liver insult and liver fibrosis regression, CD8^+^ T cell hyperfunction persisted with sustained elevated levels of cellular degranulation and IFN-γ expression, as seen in animals with ongoing liver insult.

**Figure 5:**
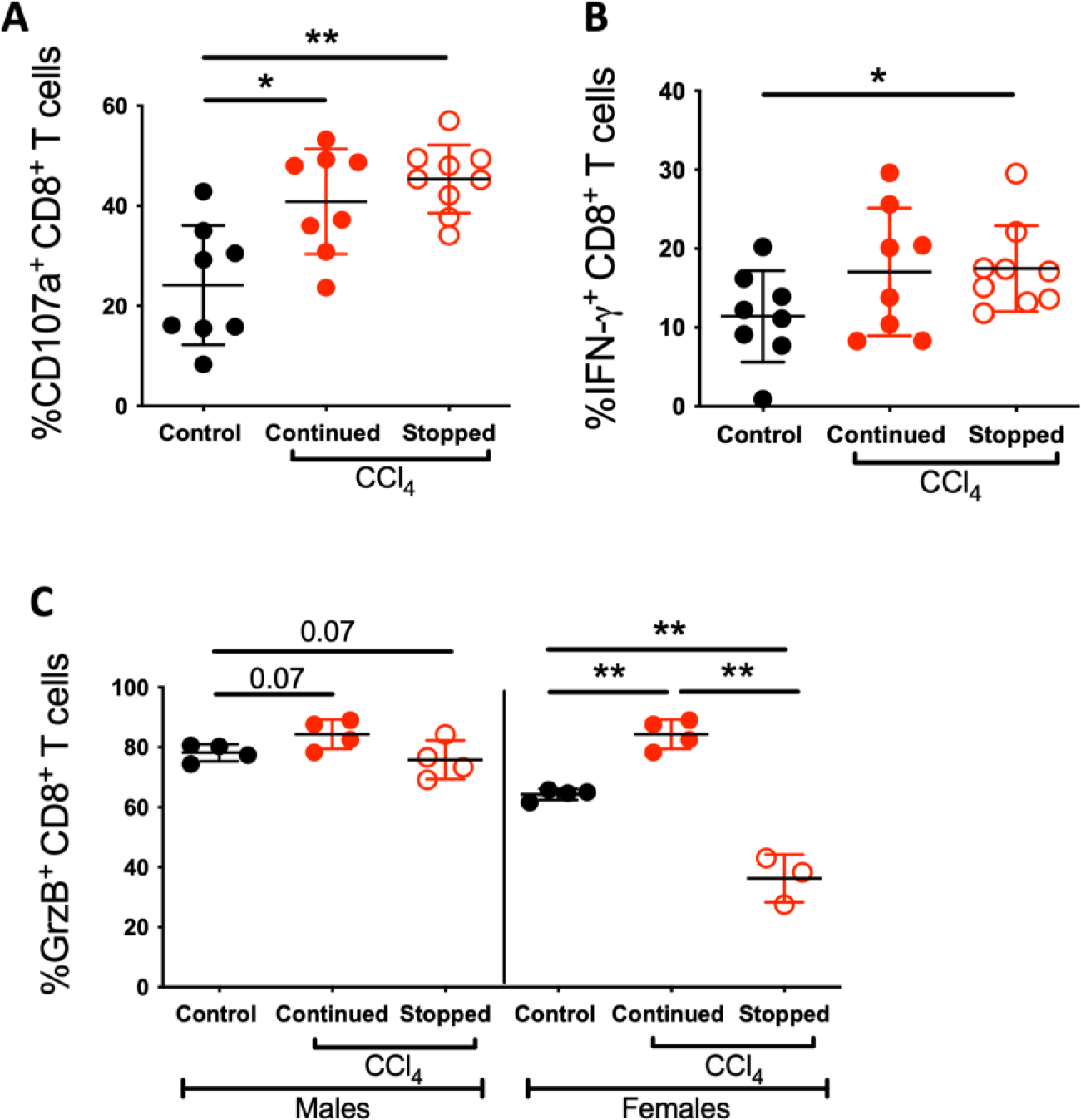
Sustained CD8^+^ T cell hyperfunction in liver fibrosis regression. CD8^+^ T cell function was assessed at 19 and 21 weeks after CCl_4_-treated mice were divided into two groups: continued CCl_4_ treatment or CCl_4_ cessation. Isolated PBMCs were stimulated *in vitro* with anti-CD3/28 for 48 hours and function was assessed by flow cytometry. The proportion of **A)** CD107a^+^ and **B)** IFN-γ^+^ CD8^+^ T cells are significantly higher in the CCl_4_-continued and cessation groups compared to controls. **C)** Notable sex differences in the proportion of GrzB^+^ CD8^+^ T cells between the former groups are shown. Data are presented as means ±SD and statistical significance was determined using a two-tailed unpaired Students’ *t*-test (*p<0.05, or **p<0.01).

### Ectopic tumour growth is increased and response to immunotherapy is delayed in mice with advanced liver fibrosis

Next, an assessment of possible clinically relevant consequences of CD8^+^ T cell hyperfunction in chronic liver fibrosis was undertaken. Following a 10–14-week course of CCl_4_ administration, known to induce CD8^+^ T cell hyperfunction, mice were challenged with tumour cells of the colon carcinoma murine cell line MC38 (Figure 6A). MC38 cells are commonly used as a transplantable tumour model^32^ and are known to induce a potent CD8^+^ T cell response^33^. When limiting doses of MC38 cells were injected (1×10^3^ or 1×10^5^ MC38 cells), control mice readily controlled tumour outgrowth, suggesting responsive anti-tumour CD8^+^ T cells. In contrast, mice with advanced liver fibrosis confirmed by histology 3 weeks post-tumour implantation, presented increasing tumour burden over time (Figure 6B and C).

**Figure 6:**
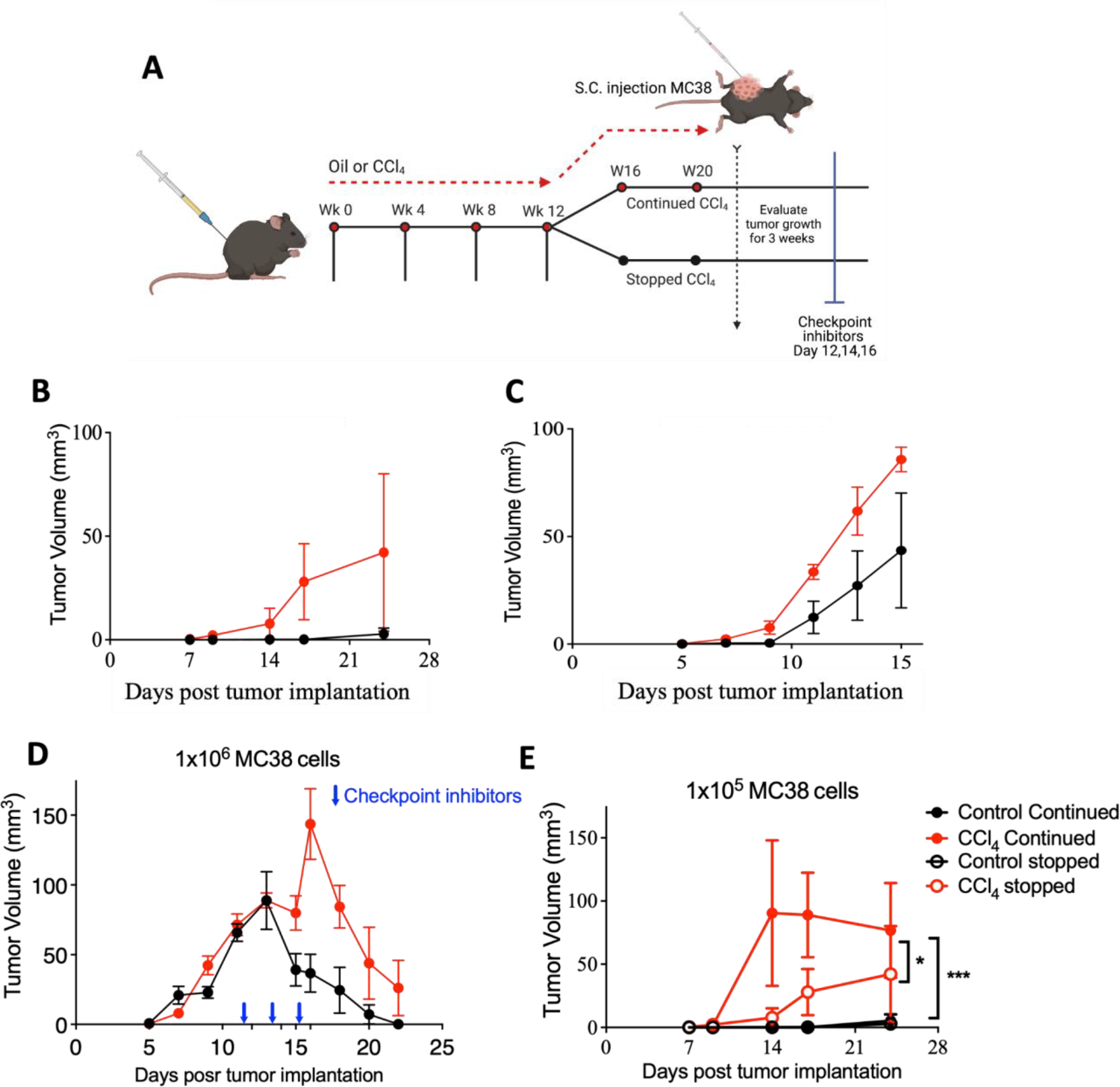
Liver fibrosis is associated with poor response to ectopic tumour challenge and delayed response to checkpoint inhibitor therapy. **A)** A schematic of the experimental design evaluating ectopic tumour challenge and response to checkpoint inhibitor therapy in a murine model of hepatotoxic liver fibrosis. Mice were injected (i.p.) with oil (control) or CCl_4_ in oil (1mL/kg body weight) twice/week for 12-16 weeks. When CD8^+^ T cell hyperfunction was detected in PBMCs *in vitro*, mice either continued to be administered CCl_4_ twice weekly or treatment was stopped. At the same time, mice were injected with MC38 tumour cells S.C. in the right flank of the abdomen to evaluate tumour growth until endpoint (≥75mm^3^). **B)** Tumour growth in control and CCl_4_-treated mice in response to 1×10^3^ MC38 cells. **C)** Oil, peak treatment (wk 12) or CCl_4_-stopped mice were then challenged with 1×10^5^ MC38 cells and tumours were measured until endpoint. Tumour growth in control and CCl_4_-treated mice in response to 1×10^5^ MC38 cells. **D)** Tumour growth in control and CCl_4_-treated mice in response to 1×10^6^ MC38, followed by injections of anti-PD1 + anti-CTLA-4 checkpoint inhibitor antibodies every two days (3 total injections). Data are presented as mean ±SD and statistical significance was determined using a two-way ANOVA to compare tumour volume in CCl_4_-treated mice vs. control mice (*p<0.05, or ***p<0.001).

To understand if liver fibrosis influences immunotherapy response, mice were injected with a higher dose of MC38 cells (1×10^6^) at peak liver fibrosis, resulting in simultaneous tumour outgrowth in both CCl_4_-treated and control mice. When tumours reached 75mm^3^, mice were treated with anti-PD-1 and anti-CTLA-4 antibodies three times over the course of 7 days. As expected, control mice promptly responded to immunotherapy, with a reduction in tumour size by day 22 p.i.. Meanwhile, their CCl_4_-treated counterparts showed a delayed response to immunotherapy (Figure 6D), failing to completely reduce tumour volume.

To determine if sustained CD8^+^ T cell hyperfunction after liver fibrosis regression is associated with a lack of tumour control, mice were subjected to a 12-week CCl_4_ treatment period and 7 weeks of regression post-CCl_4_ cessation, followed by MC38 challenge (1×10^5^ cells). Mice that received continuous liver insult developed significantly larger tumours than mice with regressed liver disease (post-cessation), while mice without fibrosis were able to fully control tumour growth (Figure 6E).

### CD8^+^ T cell hyperfunction is also associated with liver steatosis and ballooning, in the absence of significant liver fibrosis

To examine T cell function in another murine model of chronic liver insult, adult male and female C57BL/6 mice were fed a 45% HFD for 14 weeks (Figure 7A). HFD-fed mice had significantly higher body weight and percent fat mass compared to mice on a regular chow diet (Figure 7B-C, 7F-G). At week 13, diet mice were also hyperglycaemic (Figure 7D, 7H) and hyperinsulinemic (Figure 7E,7I) compared to control mice.

**Figure 7:**
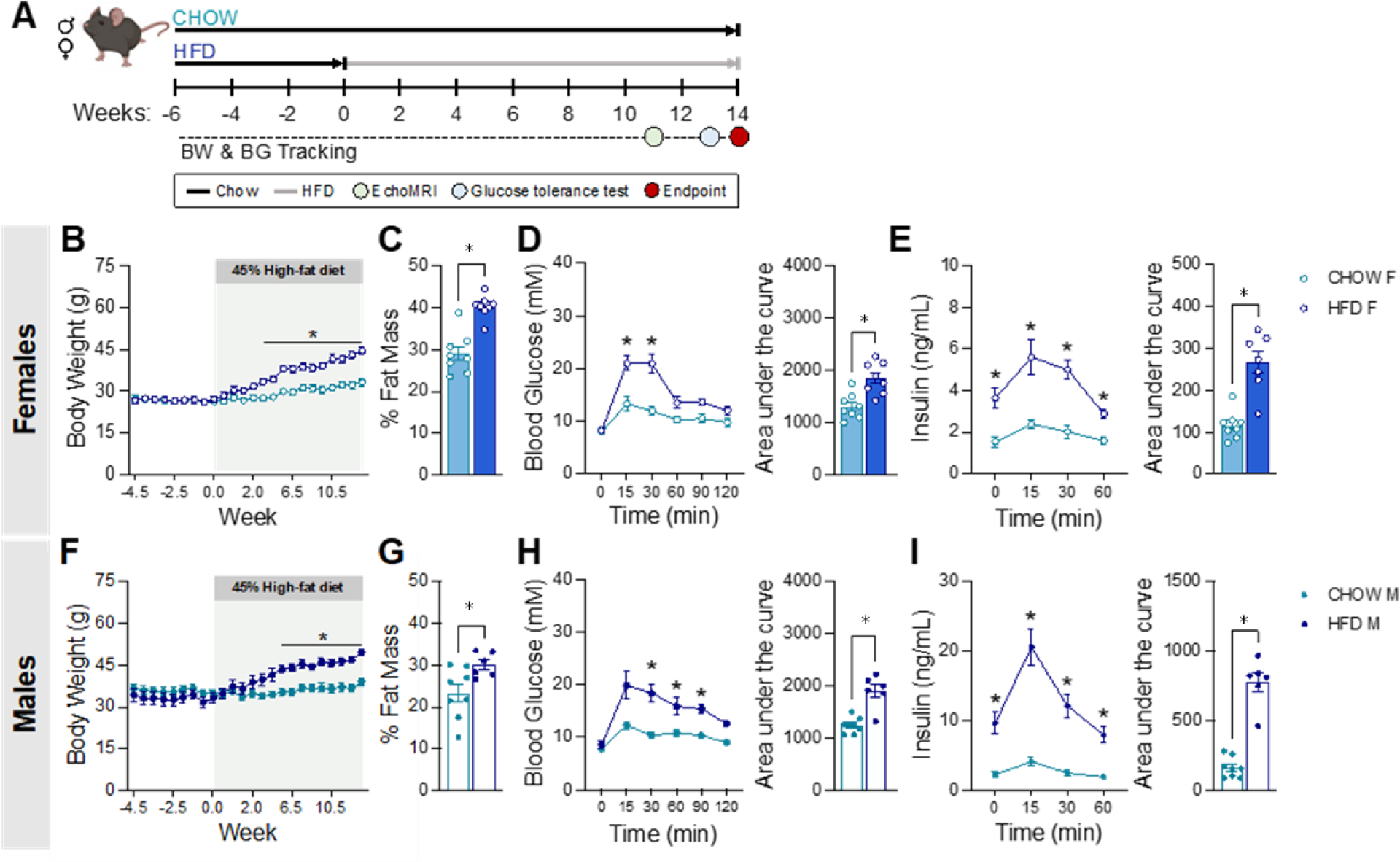
High fat diet induces liver steatosis and ballooning in the absence of significant fibrosis, following the disruption of host metabolism. **A)** Study timeline. Male and female C57BL/6 mice were kept on a regular chow diet (13% fat, 20% protein, 67% carbohydrates) after weaning. At 28 to 30 weeks of age, a subset of mice (6 males, 8 females) was transferred to a 45% HFD for 14 weeks, while the control group (8 males, 8 females) remained on the regular chow diet. **B & F)** Body weight tracking. **C & G)** Percent fat mass at week 11. **D & H)** Systemic blood glucose during the glucose tolerance test (GTT) at week 13. **E & I)** Corresponding plasma insulin levels during the GTT. Data is plotted as means ±SD and statistical significance was determined using a two-way ANOVA (*p<0.05).

Interestingly, in this model, we found that the majority of chow-fed females had developed minimal fibrosis (F1 to F2) (Figure 8A), similarly seen in chow-fed males (Figure 8C). On the other hand, in the HFD group, most males had minimal fibrosis (F0, F0.5), accompanied by low to moderate steatosis and ballooning (Figure 8D). Of these, only one mouse had stage 1 liver inflammation, and a third had low to moderate liver inflammation (stage 1 and 2). Among HFD-fed females, most had low to moderate fibrosis (F0.5 to F2), low to severe steatosis (stage 1 to 3) (Figure 8B) and stage 1 or 2 inflammation.

**Figure 8:**
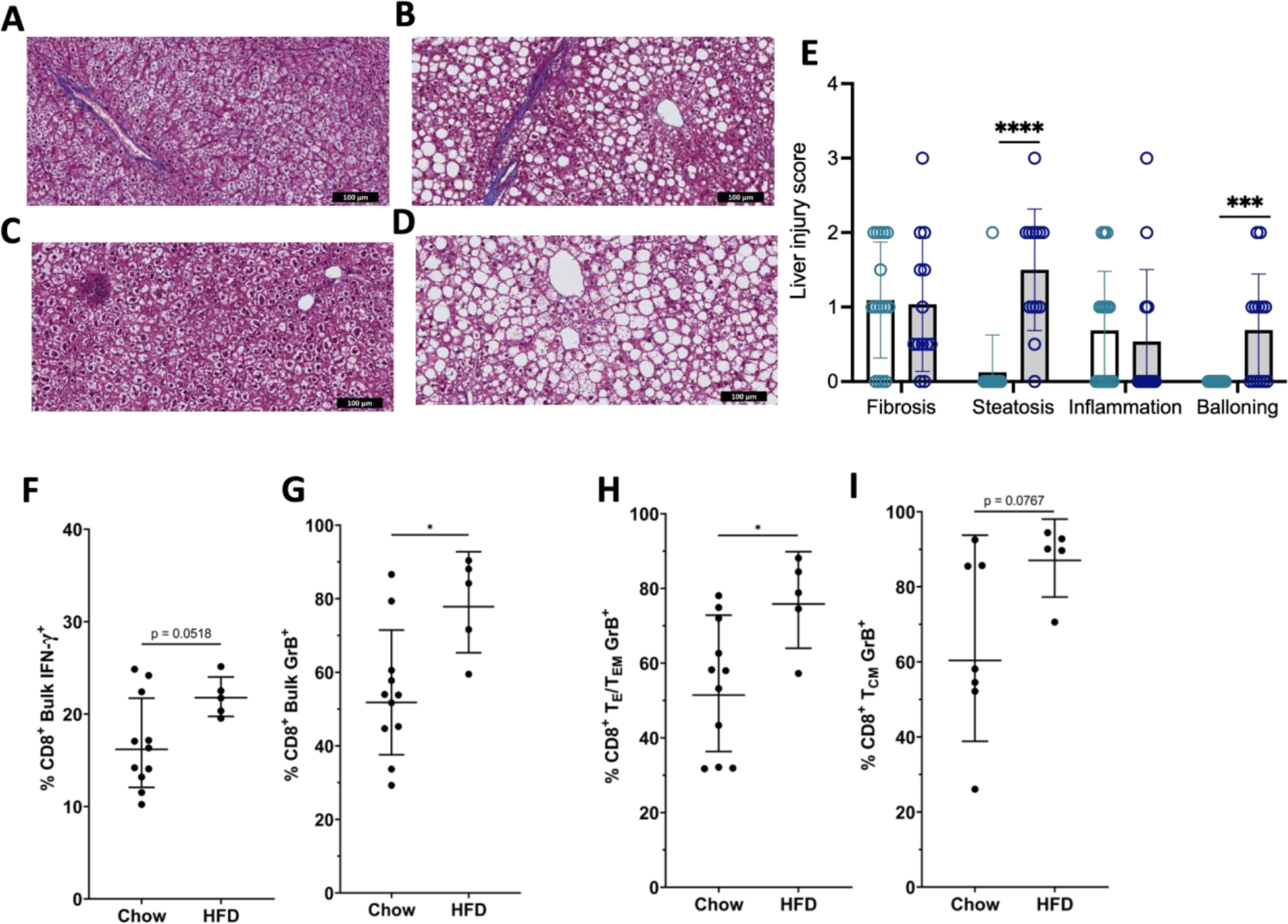
CD8^+^ T cell hyperfunction is associated with advanced liver steatosis in the presence of minimal liver fibrosis. CD8^+^ T cell function was assessed after 45% high fat diet (HFD) feeding. Male and female C57BL/6 mice were placed on a 45% HFD for 14 weeks, while the control group remained on a regular chow diet. Animals were then euthanized, and blood and liver samples collected. Liver sections were formalin-fixed, paraffin embedded, sectioned and stained with Masson’s Trichrome and haematoxylin & eosin (H&E), Isolated PBMCs were stimulated *in vitro* with anti-CD3/28 antibodies for 48 hours, and function was assessed by flow cytometry. **A-D)** Representative liver images demonstrating liver tissue pathology determined by a skilled pathologist from the Louise Pelletier Histology Core at the University of Ottawa. These tissue sections were stained with Masson’s Trichrome (fibrosis) or haematoxylin and eosin (steatosis/ballooning/inflammation): **A)** Female chow diet: fibrosis (F1), no ballooning or steatosis; **B)** Female high fat diet: fibrosis (F2), steatosis (2), no ballooning; **C)** Male chow diet: no fibrosis, no ballooning or steatosis; **D)** Male high fat diet: no fibrosis, severe steatosis (3) and ballooning (2). Complete data summary of liver pathologies in Supplemental Figure 3. **E)** Liver injury scores in chow and HFD-fed mice after the 14-week treatment period. Summary data of **F)** IFN-γ expression in bulk CD8^+^ T cells and **G)** Granzyme-B expression in bulk, **H)** T_E_/_EM_ and **I)** T_CM_ CD8^+^ T cells. Data are presented as means ±SD and statistical significance was determined using a two-tailed unpaired Students’ t-test (*p<0.05, ***p<0.001 and ****p<000.1)

HFD-fed mice have significantly elevated steatosis and ballooning in comparison to chow-fed mice (Figure 8E). Sex differences in liver injury scores are shown in supplemental figure 3. After 14 weeks of HFD feeding, we found that bulk CD8^+^ T cells in the HFD-fed group showed a significant increase in IFN-γ expression compared to the chow-fed group (Figure 8F). In addition, there was an increase in granzyme B expression in bulk cells (Figure 8G) and T_E_/T_EM_ CD8^+^ T cells (Figure 8H) as well as a trending increase in the T_CM_ subset (Figure 8I). Therefore, CD8^+^ T cell hyperfunction can be observed with chronic liver insult resulting in significant steatosis with absent or minimal fibrosis.

## Discussion

Immune cell dysfunction is a hallmark feature of chronic liver disease, associated with impaired antiviral responses to hepatic infection and increased risk of HCC. Yet, systemic immune cell hyperfunction, coupled with systemic inflammation in advanced liver damage, reported by us and others in CD8^+^ T cells, is not well understood. This study found a consistent induction of advanced liver fibrosis pathology after 12 weeks of hepatotoxin injection (i.p.), by adapting a protocol from previous studies in rats^31^. The findings reported here show that persistent presentation of severe liver pathology was associated with increased inducible expression of IFN-γ and GrzB in circulating CD8^+^ T cells. This emulates previous observations in chronic HCV infection with cirrhosis^28^. Importantly, this hyperfunction was somewhat sustained following fibrosis regression post-CCl_4_ cessation. This generalized immune dysfunction was also associated with poor control of ectopic tumour growth and a delayed response to immunotherapy. These findings not only confirmed the association of CD8^+^ T cell hyperfunction with liver fibrosis severity, but also decoupled this from hepatic viral infection. They also ascribed potential clinical relevance to long-term outcomes and therapeutic responsiveness in the context of advanced liver fibrosis. Interestingly, we complement this with our own findings showing CD8^+^ T cell hyperfunction, in a model of diet-induced chronic liver disease, causing liver steatosis without significant fibrosis. This expands the scope of impact this phenomenon holds in different etiologies of chronic liver disease.

Much has been learned about the processes of liver fibrosis progression in mouse models of liver fibrosis^22^. While there is no single animal model that recapitulates all the features of liver disease, the CCl_4_ model is characterized here by a linear development of liver fibrosis followed by periportal and portal-to-central fibrosis, both hallmark features of HCV infection pathology^22^. This data also resembles the hyperfunction patterns observed in HCV-infected individuals with cirrhosis, further confirming the suitability of this model to evaluate immune cell hyperfunction in advanced liver fibrosis. Thus, the model was well suited to decouple advanced liver fibrosis from virus, suiting our hypothesis that immune cell hyperfunction will occur in advanced liver disease of varying etiologies. The impact of fibrosis on the immune system remains relatively unknown. The CCl_4_ hepatotoxin has previously been shown to enhance then subsequently diminish mouse T cell responses to non-specific mitogens after 7 and 23 days, respectively, while impairing T cell-dependent antibody responses when administered i.p.^34, 35^. Another short-term two-week course study of CCl_4_ treatment (i.p.) resulted in impaired responses to *Listeria monocytogenes* (LCMV) and *Streptococcus pneumoniae*, both infections in which CD8^+^ T cell responses play an important role in pathogen clearance^36^. These latter studies focused on the acute effects of CCl_4_, without examining the long-term effects of hepatotoxin administration on liver pathology. To date, the model has not been evaluated in mice to study chronic advanced liver fibrosis and immunological effects.

The notion of T cell *hyperfunction* in chronic disease may be associated in part with generalized immune activation. This is widely described in chronic viral infections, characterized by increased HLA-DR and CD28 expression, as well as other biomarkers of inflammation such as serum LPS and sCD14 (Ref: PMIDs 18457674, 21726511). A recent report has found CD8^+^ T cell hyperfunction in a mouse model of non-alcoholic steatohepatitis (NASH), the inflammatory form of NAFLD, associating it with reduced tumour surveillance and poor response to immunotherapy for HCC^37^. Moreover, in this model, prophylactic immunotherapy elevated CD8^+^ T cell hyperfunction further and paradoxically increased HCC tumour burden. This sheds light on our overall hypothesis that local and systemic generalized CD8^+^ T cell hyperfunction impairs immune surveillance, a central feature of CD8^+^ T cell response, resulting in susceptibility to challenge. As it has been extensively shown that MC38 tumours face potent CD8^+^ T cell responses^33^, our data supports the notion that liver fibrosis severely compromises the immune system, to the point where it is incapable of responding to tumour challenge.

Although the removal of liver insult, such as HCV virus, can reverse or halt liver disease progression, immune dysfunction may be persistent. In HCV infection, liver fibrosis regression is often observed following sustained virologic response (i.e., virus elimination) with highly effective antiviral therapy. However, in cirrhotic individuals, occurrence of incremental regression (i.e single F-stage reduction), if at all, is ≈ 50%^29, 38^. We have observed this in previous clinical studies of HCV^+^ individuals with cirrhosis 24 weeks post-SVR with DAA therapy^39^. Nevertheless, we observed sustained CD8^+^ T cell hyperfunction long after cure^28^. Whether a lack of liver fibrosis regression is reflected in recovery from immune cell dysfunction is not known, and host or pathogen contributions therein are unclear. In a rat model of chronic CCl_4_ exposure, there was progressive loss of fibrotic matrix 28 days following the cessation of toxin administration^22,31^. We report here, for the first time in mice, that a 12-week CCl_4_ protocol achieving maximal fibrosis can subsequently be fully reversed (Figure 4). However, CD8^+^ T cell hyperfunction persisted in this time, marked by an elevated IFN-γ response (Figure 5). This suggests that removal of liver insult, or any degree of fibrosis regression, did not restore normal CD8^+^ T cell function over time. Similar retention of immune dysfunction has been reported in cirrhotic individuals when monitoring HCV-specific CD8^+^ T cell impairment post-DAA cure^16, 17, 18^. This would suggest a degree of inherent trauma to the cells, which is a challenge to overcome despite removal of the insult. This invokes the potential for lasting epigenetic modifications previously reported in other models of chronicity (Ref: PMID 27789795). Pivotal studies in the murine chronic LCMV model have revealed epigenetic modifications as an underlying feature of CD8^+^ T cell dysfunction^40^.

In addition, since any derangement of immune surveillance, an important aspect of CD8^+^ T cell function, could potentially contribute to cancer development, we also evaluated the impact on tumour challenge and immunotherapy in this model. To do so, we employed the MC38 murine cell line, a commonly used transplantable mouse tumour model^32^ known to induce a potent CD8^+^ T cell response, to investigate anticancer immunity^33^. Our data shows lack of tumour control despite the resolution of liver tissue damage (Figure 6E). These findings offer translational relevance in associating generalized systemic CD8^+^ T cell hyperfunction in advanced liver disease to impaired responses to tumour challenge. This may offer insight to the development of HCC in cirrhotic individuals, although an appropriately adapted model for liver cancer would more directly test such a hypothesis.

As there remains a calculable risk for HCC in HCV^+^ cirrhotics, despite the success of DAA therapy, this continues to be of considerable clinical significance^41^. While the risk of HCC may decrease after DAA therapy^42,43^, some studies have observed new, and often more aggressive forms of the cancer^44^, or increased recurrence^45^. These findings further highlight the value of this model in identifying targets to restore immune function, thereby enabling effective immunotherapies for cancer. The data may provide insights on why liver cancer patients, where fibrosis is often present, may respond sub-optimally to PD-1 or CTLA-4 immunotherapy^46, 47^.

These results were complemented by a study of bulk CD8^+^ T cell function in a high fat diet (HFD) model of induced chronic liver damage. The 45% HFD better recapitulates liver pathology, metabolic syndrome and disease pathogenesis observed in human NAFLD, in comparison to the previously used hepatotoxin model. This HFD induces little to no liver fibrosis, decoupling it from advanced steatosis (Figure 7K and M). This is achieved by inducing glucose intolerance, subsequently resulting in increased fat mass and body weight (Figure 7A-I). This liver insult was associated with statistically significant increases in CD8^+^ T cell function (Figure 8), resembling that observed in the CCl_4_ model. In a 60% HFD study, it was found that, after 12 weeks of diet feeding, mice had elevated Th17/T_reg_ ratios in adipose tissue, increasing serum proinflammatory cytokines^48^. It was found that T cells are known to regulate chronic inflammation while contributing to abnormal energy metabolism^49^. While the relationship between obesity, chronic inflammation, and metabolic syndrome remains unclear, our findings implicate the phenomenon of CD8^+^ T cell hyperfunction in metabolic syndrome affecting the liver. Given the common immune dysregulation in these models of chronic liver injury, clinical outcomes in cancer and immunotherapy may be compromised in steatotic liver disease.

The burden of chronic liver disease on the health of millions, either of infectious or non-infectious etiologies, is mounting. Over 50 million people are affected by chronic HCV infection worldwide, of which ≈15-30% will silently develop cirrhosis and symptomatic liver disease, predisposing to end-stage liver disease and high mortality rates^50, 51^. It is predicted that half of HCV^+^ individuals in the USA will develop cirrhosis by 2030, often occurring before diagnosis or treatment, indicating that the relevancy of research on the impact of liver damage on the host will remain significant for decades^52^. In addition, the rising burden of inflammatory fatty liver disease presents a growing challenge to global public health. The use of mouse models emulating the generalized CD8^+^ T cell dysfunction and liver fibrosis observed in humans^28^, will enable mechanistic investigations, and offer understanding of the impact on clinical outcomes and potential for therapeutic efficacy.

## Acknowledgements

This research has been funded by the Canadian Institutes of Health Research (CIHR) Operating Grant (PJT-419982) to Crawley and a National Sciences and Engineering Research Council of Canada (NSERC) Discovery Grant (#RGPIN-2017-06265) to Bruin. Madani J was supported by a stipend from the Saudi Arabian government; Li J is funded by a Canadian Network on Hepatitis C Graduate Scholarship, and a Queen Elizabeth II Graduate Scholarship in Science and Technology from the Government of Ontario; Jorritsma K is funded by scholarships from the Center for Infection, Immunity and Inflammation (University of Ottawa). Jorritsma J and Ching AC are both funded by the Canadian Graduate Scholarships.

We gratefully acknowledge the services provided by the Louise Pelletier Histology Core Facility (RRID: SCR_021737) at the University of Ottawa for mouse liver tissue embedding, processing, staining, and imaging. We also gratefully acknowledge the services provided by the Flow Cytometry and Cell Sorting Core Facility (RRID: SCR_023349) at the Ottawa Hospital Research Institute, and the Flow Cytometry and Virometry Core Facility (RRID: SCR_023306) at the University of Ottawa, for access to high-throughput multi-parameter flow cytometry tools and equipment.

We would also like to thank Natasha Campeau (Crawley AM lab) for assisting in minor edits and providing valuable feedback during the writing of the manuscript.

## Author contributions

JM performed many of the CCl_4_ experiments, analyzed the data and contributed to the writing of the manuscript. JL assisted with some of the CCl_4_ experiments. AV performed some of the CCl_4_ experiments, trained staff and assisted in supervising the laboratory experiments. JL and KJ conducted the HFD T-cell experiments and analysis and aided in editing the manuscript. MSH assisted in the first CCl_4_ experiment and tumour measurements. MD performed the pathology scoring of tissue sections. ACC provided important clinical direction to inform study design, data interpretation and editing of the manuscript. AC conducted the HFD experiments and related metabolic analysis and contributed to the writing of the manuscript. JEB designed the HFD experiments, assisted in data analysis, and contributed to the writing and editing of the manuscript. MA aided in the design of the CCl_4_ studies, assisted with data analysis and contributed to the writing and editing of the manuscript. AMC designed the CCl_4_ research project, supervised laboratory experiments, assisted with data analysis, brokered collaboration with JEB team and wrote and edited the manuscript.

## Figure legends

**Supplemental Figure 1: Weight gain does not differ between control and CCl_4_-treated mice.** Mice were administered 200 µL of PBS or CCl_4_ (0.5 ml/kg) dissolved in olive oil (50% volume) twice weekly, by i.p. injections. A representative data set of weekly recorded body weights in ≈ 7-week-old C57BL/6 mice, who were administered oil (controls) or CCl_4_, i.p. for 12 weeks.

**Supplemental Figure 2: Detection of elevated CD107a expression in blood CD8^+^ T cells is associated with advanced liver fibrosis.** Isolated PBMCs were stimulated *in vitro* with anti-CD3/28 antibodies for 48 hours, and function was assessed by flow cytometry. Summary data of **A)** CD107a expression in control mice and CCl_4_-treated mice after 12 weeks of the treatment. **B)** CD107a expression was evaluated in CD8^+^ T cell subsets by flow cytometry using the following subset distinguishing phenotypes: naïve (CD44^low^CD62L^+^), effector (CD44^+^CD62L^-^), and memory (CD44^+^CD62L^+^). Data is plotted as means ±SD and statistical significance was determined using a two-tailed unpaired Students’ *t*-test (*p<0.05).

**Supplemental Figure 3: Sex differences in high-fat diet-induced liver damage.** Male and female C57BL/6 mice were placed on a regular chow diet (13% fat, 20% protein, 67% carbohydrates) after weaning. At 28 to 30 weeks of age, a subset of mice (6 males, 8 females) was transferred to a 45% HFD for 14 weeks, while the control group (8 males, 8 females) remained on a regular chow diet. Animals were then euthanized, and livers were harvested, formalin-fixed, and paraffin embedded, sectioned, and stained with Masson’s Trichrome three colour stain. Liver tissue damage was assessed by a pathologist. Liver fibrosis severity is summarized in **A)** males and **B)** females. Fibrosis staging was determined using the METAVIR (F-score) scoring system: F0 = no fibrosis, F1/2 = minimal liver fibrosis, F3/4 = advanced fibrosis. Degree of steatosis, tissue ballooning, and inflammatory staging were also determined.

## Notes

### Competing Interest Statement

The authors have declared no competing interest.

## References

1. Noor, M.T. & Manoria, P. Immune Dysfunction in Cirrhosis. J Clin Transl Hepatol 5, 50–58 (2017).

2. Albillos, A., Lario, M. & Alvarez-Mon, M. Cirrhosis-associated immune dysfunction: distinctive features and clinical relevance. J Hepatol 61, 1385–1396 (2014).

3. Swain, M.G. et al. Burden of nonalcoholic fatty liver disease in Canada, 2019-2030: a modelling study. CMAJ Open 8, E429–E436 (2020).

4. Penna, A. et al. Dysfunction and functional restoration of HCV-specific CD8 responses in chronic hepatitis C virus infection. Hepatology 45, 588–601 (2007).

5. Gruener, N.H. et al. Sustained dysfunction of antiviral CD8+ T lymphocytes after infection with hepatitis C virus. J Virol 75, 5550–5558 (2001).

6. Rehermann, B. Chronic infections with hepatotropic viruses: mechanisms of impairment of cellular immune responses. Semin Liver Dis 27, 152–160 (2007).

7. Cox, A.L. et al. Comprehensive analyses of CD8+ T cell responses during longitudinal study of acute human hepatitis C. Hepatology 42, 104–112 (2005).

8. Wedemeyer, H. et al. Impaired effector function of hepatitis C virus-specific CD8+ T cells in chronic hepatitis C virus infection. J Immunol 169, 3447–3458 (2002).

9. Shen, T., Chen, X., Xu, Q., Lu, F. & Liu, S. Distributional characteristics of CD25 and CD127 on CD4+ T cell subsets in chronic HCV infection. Arch Virol.

10. Spaan, M. et al. Immunological Analysis During Interferon-Free Therapy for Chronic Hepatitis C Virus Infection Reveals Modulation of the Natural Killer Cell Compartment. J Infect Dis 213, 216–223 (2016).

11. Hengst, J. et al. Nonreversible MAIT cell-dysfunction in chronic hepatitis C virus infection despite successful interferon-free therapy. Eur J Immunol 46, 2204–2210 (2016).

12. Sumida, K. et al. Characteristics of splenic CD8+ T cell exhaustion in patients with hepatitis C. Clin Exp Immunol 174, 172–178 (2013).

13. Rehermann, B. et al. Quantitative analysis of the peripheral blood cytotoxic T lymphocyte response in patients with chronic hepatitis C virus infection. J Clin Invest 98, 1432–1440 (1996).

14. Zhao, B.B. et al. T lymphocytes from chronic HCV-infected patients are primed for activation-induced apoptosis and express unique pro-apoptotic gene signature. PLoS One 8, e77008 (2013).

15. Lucas, M. et al. Pervasive influence of hepatitis C virus on the phenotype of antiviral CD8+ T cells. J Immunol 172, 1744–1753 (2004).

16. Osuch, S., Metzner, K.J. & Caraballo Cortes, K. Reversal of T Cell Exhaustion in Chronic HCV Infection. Viruses 12 (2020).

17. Owusu Sekyere, S. et al. Type I Interferon Elevates Co-Regulatory Receptor Expression on CMV- and EBV-Specific CD8 T Cells in Chronic Hepatitis C. Front Immunol 6, 270 (2015).

18. Perpinan, E. et al. Chronic genotype 1 hepatitis C along with cirrhosis drives a persistent imprint in virus-specific CD8(+) T cells after direct-acting antiviral therapies. J Viral Hepat 27, 1408–1418 (2020).

19. Burke Schinkel, S.C., Carrasco-Medina, L., Cooper, C.L. & Crawley, A.M. Generalized Liver- and Blood-Derived CD8+ T-Cell Impairment in Response to Cytokines in Chronic Hepatitis C Virus Infection. PLoS One 11, e0157055 (2016).

20. Vranjkovic A., D.F., Kaka S., Cooper CL., Crawley AM. Dysfunction of circulating CD8+ T-cells in chronic HCV infection is associated with severity of liver disease and is unresolved after HCV cure. Frontiers in Immunology In press (2019).

21. Moreno-Cubero, E. et al. According to Hepatitis C Virus (HCV) Infection Stage, Interleukin-7 Plus 4-1BB Triggering Alone or Combined with PD-1 Blockade Increases TRAF1(low) HCV-Specific CD8(+) Cell Reactivity. J Virol 92 (2018).

22. Delire, B., Starkel, P. & Leclercq, I. Animal Models for Fibrotic Liver Diseases: What We Have, What We Need, and What Is under Development. J Clin Transl Hepatol 3, 53–66 (2015).

23. Constandinou, C., Henderson, N. & Iredale, J.P. Modeling liver fibrosis in rodents. Methods Mol Med 117, 237–250 (2005).

24. Kofahi, H.M., Taylor, N.G., Hirasawa, K., Grant, M.D. & Russell, R.S. Hepatitis C Virus Infection of Cultured Human Hepatoma Cells Causes Apoptosis and Pyroptosis in Both Infected and Bystander Cells. Sci Rep 6, 37433 (2016).

25. Domenicali, M. et al. A novel model of CCl4-induced cirrhosis with ascites in the mouse. J Hepatol 51, 991–999 (2009).

26. Hillebrandt, S., Goos, C., Matern, S. & Lammert, F. Genome-wide analysis of hepatic fibrosis in inbred mice identifies the susceptibility locus Hfib1 on chromosome 15. Gastroenterology 123, 2041–2051 (2002).

27. Xie, C., Ma, B., Wang, N. & Wan, L. Comparison of serological assessments in the diagnosis of liver fibrosis in bile duct ligation mice. Exp Biol Med (Maywood) 242, 1398–1404 (2017).

28. Vranjkovic, A. et al. Direct-Acting Antiviral Treatment of HCV Infection Does Not Resolve the Dysfunction of Circulating CD8(+) T-Cells in Advanced Liver Disease. Front Immunol 10, 1926 (2019).

29. Casado, J.L. et al. Regression of liver fibrosis is progressive after sustained virological response to HCV therapy in patients with hepatitis C and HIV coinfection. J Viral Hepat 20, 829–837 (2013).

30. Rockey, D.C. & Friedman, S.L. Fibrosis Regression After Eradication of Hepatitis C Virus: From Bench to Bedside. Gastroenterology 160, 1502–1520 e1501 (2021).

31. Iredale, J.P. et al. Mechanisms of spontaneous resolution of rat liver fibrosis. Hepatic stellate cell apoptosis and reduced hepatic expression of metalloproteinase inhibitors. J Clin Invest 102, 538–549 (1998).

32. Corbett, T.H., Griswold, D.P., Jr., Roberts, B.J., Peckham, J.C. & Schabel, F.M., Jr. Tumor induction relationships in development of transplantable cancers of the colon in mice for chemotherapy assays, with a note on carcinogen structure. Cancer Res 35, 2434–2439 (1975).

33. Enamorado, M. et al. Enhanced anti-tumour immunity requires the interplay between resident and circulating memory CD8(+) T cells. Nat Commun 8, 16073 (2017).

34. Jirova, D., Sperlingova, I., Halaskova, M., Bendova, H. & Dabrowska, L. Immunotoxic effects of carbon tetrachloride--the effect on morphology and function of the immune system in mice. Cent Eur J Public Health 4, 16–20 (1996).

35. Delaney, B., Strom, S.C., Collins, S. & Kaminski, N.E. Carbon tetrachloride suppresses T-cell-dependent immune responses by induction of transforming growth factor-beta 1. Toxicol Appl Pharmacol 126, 98–107 (1994).

36. Guo, T.L. et al. Carbon tetrachloride is immunosuppressive and decreases host resistance to Listeria monocytogenes and Streptococcus pneumoniae in female B6C3F1 mice. Toxicology 154, 85–101 (2000).

37. Pfister, D. et al. NASH limits anti-tumour surveillance in immunotherapy-treated HCC. Nature (2021).

38. Shiratori, Y. et al. Histologic improvement of fibrosis in patients with hepatitis C who have sustained response to interferon therapy. Ann Intern Med 132, 517–524 (2000).

39. Doyle M-A., G.C., Mulvihill E., Crawley, AM., Cooper CL. Hepatitis C Direct Acting Antivirals and Ribavirin Modify Lipid but not Glucose Parameters. Cells (Revisions submitted, Feb. 2019).

40. Sen, D.R. et al. The epigenetic landscape of T cell exhaustion. Science 354, 1165–1169 (2016).

41. Lockart, I., Yeo, M.G.H., Hajarizadeh, B., Dore, G.J. & Danta, M. HCC incidence after hepatitis C cure among patients with advanced fibrosis or cirrhosis: A meta-analysis. Hepatology 76, 139–154 (2022).

42. Alberti, A. & Piovesan, S. Increased incidence of liver cancer after successful DAA treatment of chronic hepatitis C: Fact or fiction? Liver Int 37, 802–808 (2017).

43. Romano, A. et al. Newly diagnosed hepatocellular carcinoma in patients with advanced hepatitis C treated with DAAs: A prospective population study. J Hepatol 69, 345–352 (2018).

44. Romano, A., Franco Capra, Sara Piovesan, Liliana Chemello, Luisa Cavalletto, Georgios Anastassopoulos, Valter Vincenzi, PierGirogio Scotton, Sandro Panese, Diego Tempesta, Martina Gambato, Francesco P. Russo, Tosca Bertin, Maurizio Carrara, Antonio Carlotto, Giada Carolo, Giovanna Scroccaro, Alfredo Alberti. Incidence and pattern of “de novo” hepatocellular carcinoma in HCV patients treated with oral DAAs. American Association of Liver Diseases, 2018 Conference. (2018).

45. Reig, M., Boix, L. & Bruix, J. The impact of direct antiviral agents on the development and recurrence of hepatocellular carcinoma. Liver Int 37 Suppl 1, 136–139 (2017).

46. Heinrich, B., Czauderna, C. & Marquardt, J.U. Immunotherapy of Hepatocellular Carcinoma. Oncol Res Treat 41, 292–297 (2018).

47. Liu, X. & Qin, S. Immune Checkpoint Inhibitors in Hepatocellular Carcinoma: Opportunities and Challenges. Oncologist 24, S3–S10 (2019).

48. Kiran, S., Rakib, A., Kodidela, S., Kumar, S. & Singh, U.P. High-Fat Diet-Induced Dysregulation of Immune Cells Correlates with Macrophage Phenotypes and Chronic Inflammation in Adipose Tissue. Cells 11 (2022).

49. Surendar, J. et al. Adiponectin Limits IFN-gamma and IL-17 Producing CD4 T Cells in Obesity by Restraining Cell Intrinsic Glycolysis. Front Immunol 10, 2555 (2019).

50. Polaris Observatory, H.C.V.C. Global prevalence and genotype distribution of hepatitis C virus infection in 2015: a modelling study. Lancet Gastroenterol Hepatol 2, 161–176 (2017).

51. Seeff, L.B. Natural history of chronic hepatitis C. Hepatology 36, S35–46 (2002).

52. Davis, G.L., Alter, M.J., El-Serag, H., Poynard, T. & Jennings, L.W. Aging of hepatitis C virus (HCV)-infected persons in the United States: a multiple cohort model of HCV prevalence and disease progression. Gastroenterology 138, 513–521, 521 e511-516 (2010).

