## Supplemental figures for "CD8 T cell hyperfunction and reduced tumour control in models of advanced liver fibrosis"

**Supplemental Figure 1**

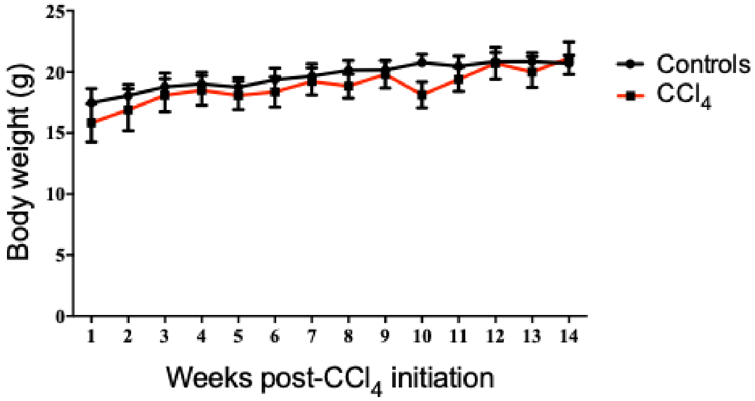


**Supplemental Figure 2**

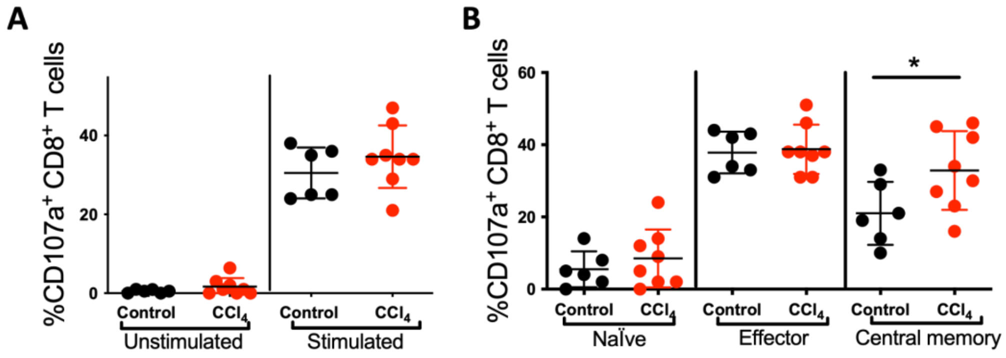


**Supplemental Figure 3**

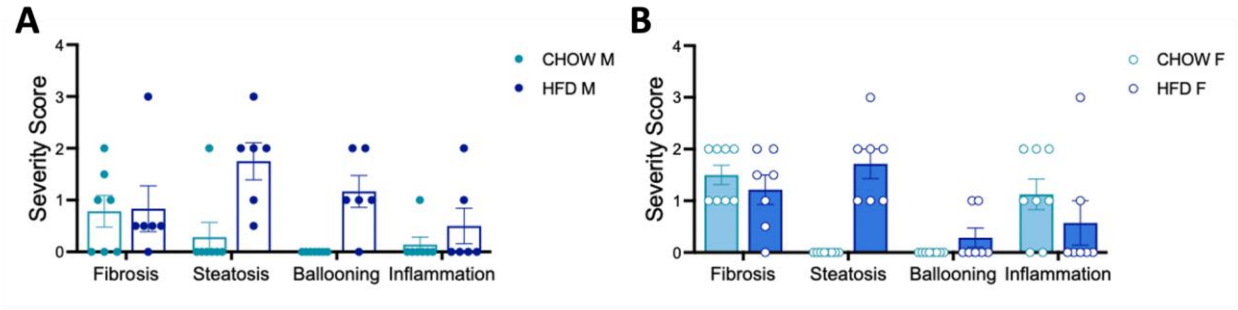
